# Development and characterisation of pNarsenic: a naringenin-inducible biosensor for arsenic in *Escherichia coli*

**DOI:** 10.1101/2025.09.25.678503

**Authors:** Vincent Crabbe, Ezgi Unal, Stijn De Graeve, Daniel G. Guerra, Tom Peeters, Sophie de Buyl, Eveline Peeters, Indra Bervoets

## Abstract

Whole-cell biosensors detecting the heavy metal arsenic have been widely studied for their potential in environmental monitoring. And while inducible biosensors have been shown to be an effective tool to tune the operational range, a thoroughly characterised inducible biosensor is currently lacking. Here, we present an *Escherichia coli* biosensor for arsenic in which the transcription factor gene *arsR* is inducible by naringenin, a plant-derived secondary metabolite. Increasing naringenin concentrations reduced the basal output while increasing both the dynamic range and sensing threshold of the biosensor dose-response curves, but the operational ranges appeared constrained by a fixed upper limit. Comparison with a previously published phenomenological model revealed good overall agreement between experimental data and model predictions, except for the behaviour of the maximum output and threshold. This work expands the biosensor toolbox with a profoundly characterised arsenic biosensor and raises a potential practical limit to dose-response curve engineering by tuning transcription factor expression alone.

**Graphical abstract:** 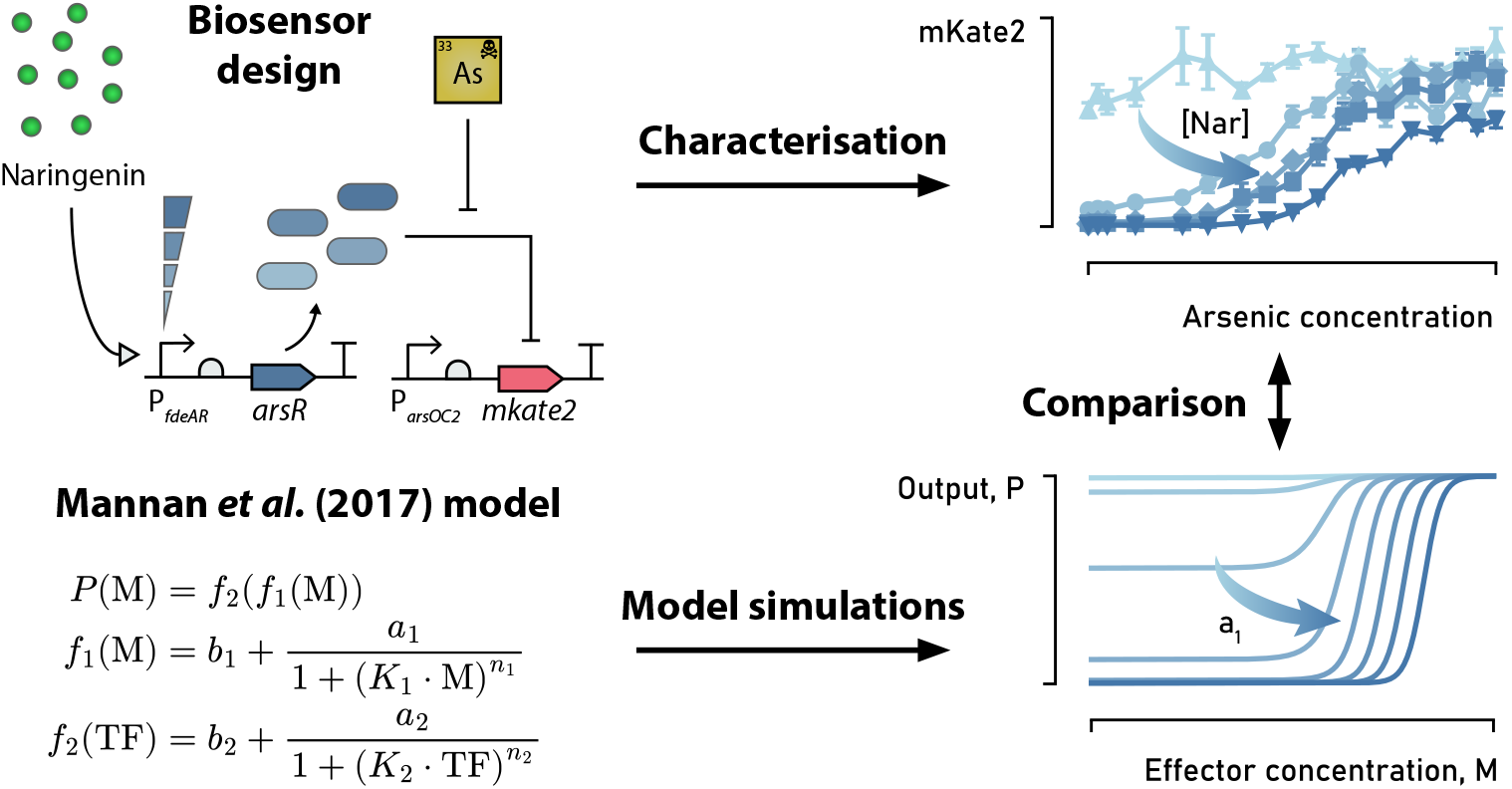

## 1 Introduction

Whole-cell biosensors have been extensively studied for environmental monitoring as they are cost-effective, sensitive, specific, and self-renewing [1, 2]. Among these, arsenic biosensors have received particular attention due to the widespread contamination of groundwater with arsenic [3] and the availability of the well-characterised transcription factor (TF) ArsR from *E. coli* [4].

ArsR functions as a repressed-repressor. In the absence of its effector, the ArsR dimer binds to its TF binding site (TFBS) upstream of the P_*ars*_ promoter, blocking transcription of the output gene. Binding to its effector arsenic causes its dissociation from the TFBS allowing transcription to proceed [5, 6]. The whole of these interactions govern the shape of the dose-response curve. Often, the native dose-response curve of a biosensor does not match the intended application, necessitating response curve engineering [7].

Because of the complexity of biosensors, synthetic biologists have used mathematical modelling to guide response curve engineering. The phenomenological model for TF-based biosensors of Mannan *et al*. [8] has gained a lot of attention by the biosensor community. It predicts the biosensor output based on eight parameters. The model makes minimal mechanistic assumptions, and should therefore be applicable across diverse mechanisms and TF families. The original publication focused on the effect of tuning the affinity of the TF to its TFBS and to its effector as well as the strength of the promoter controlling the output gene, and compared it to experimental data of LacI-based biosensors. Despite the attention their work has received, to our knowledge, their work has not yet been extended to other biosensors or to the other parameters.

Previous studies have shown that the expression level of *arsR* strongly affects the response characteristics of the biosensor. Mathematical modelling of genetic circuits showed that optimal sensitivity can be attained by placing *arsR* under control of a weak promoter while uncoupling its native feedback loop [9]. Wang and colleagues (2015) tested four biosensors with *arsR* expressed from constitutive promoters of different strength. They found that the sensitivity and output fluorescence decreased for higher *arsR* expression levels [7]. In both studies, biosensors with inducible expression of *arsR* were developed as well, but thorough characterisation including fitting to the Hill function and delineation of the operational range were omitted, making it impractical for the synthetic biology community to reuse them.

In this study, we developed a novel biosensor module for arsenic named *pNarsenic*. pNarsenic harbours *arsR* under control of the tight *fdeR*-P_*fdeAR*_ naringenin-inducible system [10]. We measured detailed dose-response curves in *E. coli* in the presence of 4 different naringenin concentrations and report fits to a Hill function and operational ranges. Besides, we compared our results with simulations of the phenomenological model of Mannan and colleagues [8] for perturbations in *a*_1_, the maximum increase in TF expression. Our study expands the analysis of the phenomenological model of Mannan *et al*. to the TF expression level, a crucial parameter for biosensor development. Further, we expanded the biosensor toolbox with the plug-and-play development of a new arsenic biosensor.

## 2 Materials and methods

Additional experimental details are provided in the Supplementary Information.

### 2.1 Reporter gene assays and data analysis

Precultures were started by inoculating separate, freshly transformed colonies in 200 µL of MOPS EZ Rich Defined Medium in a 96-well plate and incubating them overnight at 30°C and 400 rpm. Next, naringenin dissolved in ethanol was added to the desired concentration in a black 96-well plate (Greiner Bio-One, polystyrene, F-bottom), after which the ethanol was evaporated at 50°C for 1 h in sterile conditions. Next, 170 µL of MOPS EZ Rich Defined Medium was added to each well and 20 µL of the appropriate stock solution of NaAsO_2_ was added to create an arsenic concentration gradient. Finally, 10 µL of 15X diluted preculture was added, after which the plate was sealed using a Breathe-Easy Sealing Membrane.

Plates were incubated at 30°C and 425 rpm in a Biotek Synergy H1 microplate reader, measuring OD_600_ and mKate2 fluorescence (excitation: 588 nm, emission: 633 nm, gain: 110, bottom read) every 20 min. Four biological replicates were analysed per biosensor strain and arsenic concentration.

Reporter gene expression was analysed as previously described [11] with slight modifications. Briefly, medium without cell culture was used to correct for background optical density (OD_med_) and fluorescence (FL_med_). Considering three subsequent time points around 12 h of incubation (indicated as *t*\*1, *t, t*+1), the normalised corrected fluorescence for each replicate could be calculated as follows:

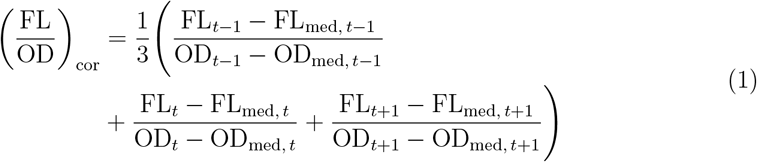

Replicates identified as outliers were entirely excluded from the dataset to avoid bias in curve fitting. The resulting response curve was fitted to a Hill function of the shape

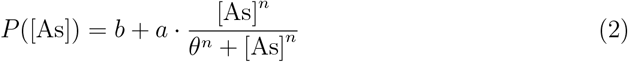

with *P* representing the normalised corrected fluorescence as calculated from (1), [As] the concentration of NaAsO_2_ in the growth medium in µM, *b* the basal output, *a* the maximum increase in output, *θ* the sensing threshold and *n* the Hill coefficient. The dynamic range *µ* was calculated as

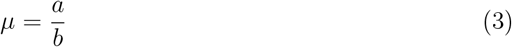

The operational range OR was computed as

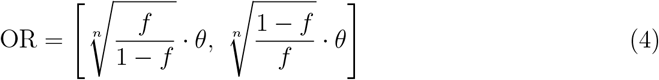

with *f* a fraction fixed at 0.1 for this article, and *θ* and *n* the best-fit threshold value and Hill coefficient for that response curve. Refer to Supplementary Information for the derivation of this formula.

The Noise parameter was defined as the average relative standard deviation

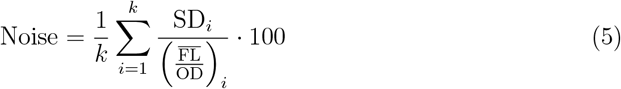

where *k* is the number of arsenic concentration levels and SD_*i*_ and 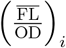, represent the standard deviation and mean normalised corrected fluorescence, respectively, at arsenic concentration *i*.

### 2.2 Mathematical modelling

We considered the repressed-repressor architecture of the phenomenological model of Mannan *et al*. for biosensor response [8]. The model computes the response *P* as a composite function

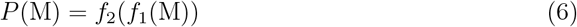

where *f*_1_ represents the concentration of “active” repressor that can bind to its TFBS as a function of the effector concentration M and *f*_2_ represents the expression level of the output gene *P* as a function of the active repressor concentration TF:

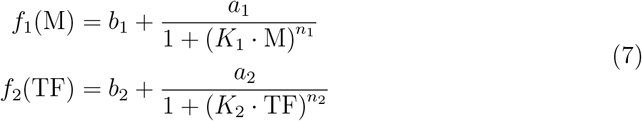

For brevity, the parameter definitions and formulae for the basal output, maximum increase in output, threshold and dynamic range can be consulted in the Supplementary Information (Table S1 and equations (8) to (11)). All computations were performed in Python; SymPy was used for symbolic computations.

## 3 Results and discussion

### 3.1 Construction of pNarsenic

To precisely control the expression level of *arsR*, we cloned the chromosomal *arsR* gene from *E. coli* K-12 downstream of the *fdeR*-P_*fdeAR*_ naringenin-inducible system [10] (Figure 1A). FdeR is a TF from *Herbaspirillum seropedicae* that functions as a dual-function regulator in *E. coli* : it represses the P_*fdeA*_ promoter in the absence of the plant secondary metabolite naringenin and induces it in the presence of naringenin [12, 13]. We chose this system because it displays an analogue response to naringenin with very tight regulation and naringenin shows low toxicity to *E. coli* [10, 13]. Further, we chose mKate2 as reporter because of its fast maturation, high pH resistance and far-red emission spectrum which limits interference with autofluorescence emitted by *E. coli* [10, 14]. We cloned *mkate2* under transcriptional control of P_*arsOC2*_, a refactored version of P_*ars*_ with lower leakiness and improved dynamic range [15]. These circuits were combined with a high-copy pUC origin of replication, yielding the 5.2 kB plasmid pNarsenic (Figure 1A).

**Figure 1:**
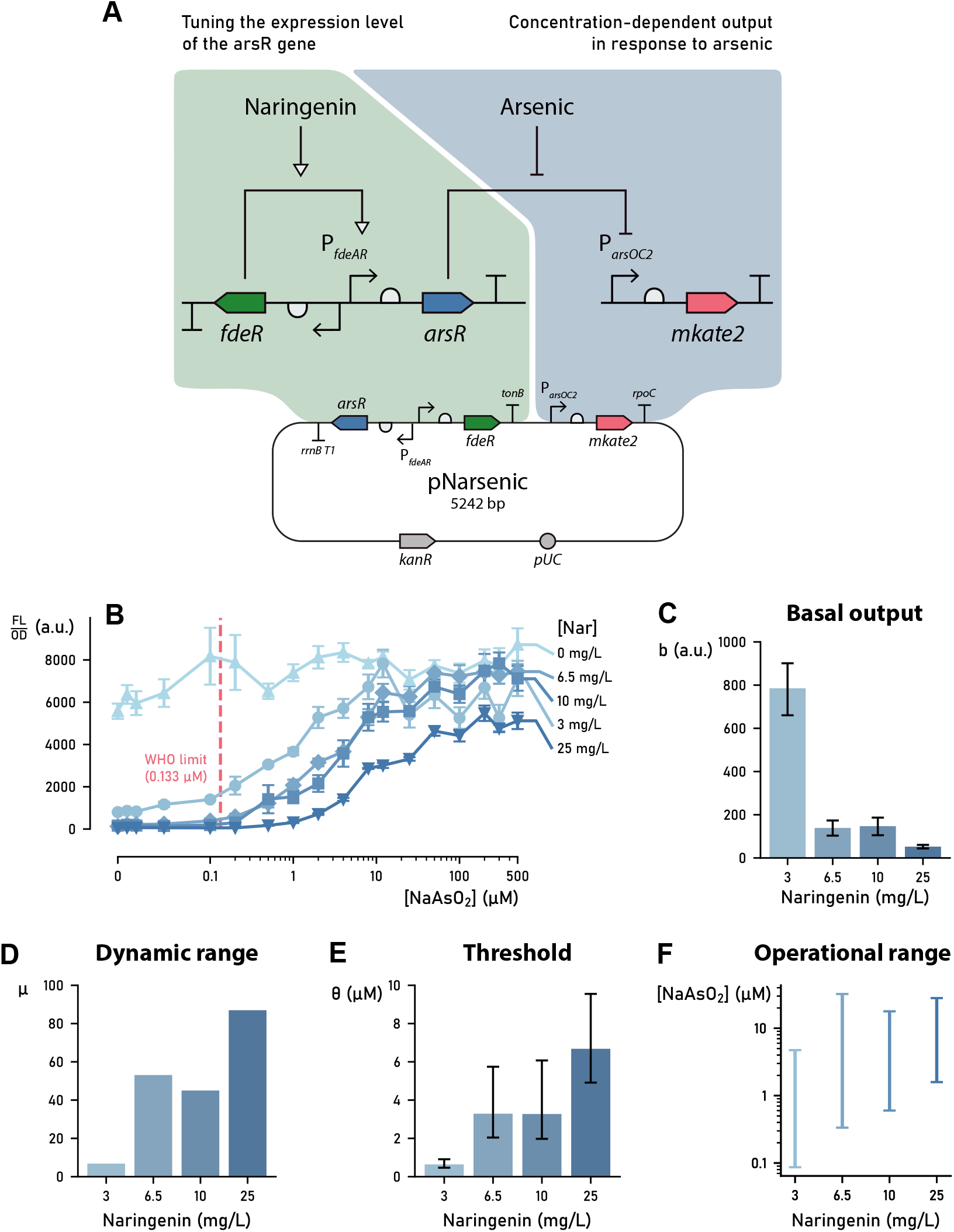
The response characteristics of the pNarsenic biosensor can be tuned by supplementing the flavonoid naringenin. (A) Plasmid map of pNarsenic with close-up of the most important features. Expression of *arsR* is induced by naringenin and repression of P_*arsOC2*_ by ArsR is relieved by arsenic, leading to mKate2 expression. (B) Dose-response curves of the pNarsenic biosensor in the presence of 0, 3, 6.5, 10, or 25 mg/L naringenin. The x-axis scale is linear between 0 µM and 0.1 µM and logarithmic between 0.1 µM and 500 µM. Error bars signify mean ± standard error across three replicates, except for 0 mg/L naringenin where *n* = 4. Refer to Figure S1 to view the response curves separately, along with their best-fit Hill functions. (C-E) Bar plots showing the basal output *b*, dynamic range *µ* and threshold *θ* for the response curves in panel B. Error bars signify the 95% confidence intervals after nonlinear regression; note that these are not available for the dynamic ranges as they were computed from equation (3). (G) The operational ranges of the response curves shown in panel B as computed through equation (4).

### 3.2 In vivo characterisation of pNarsenic

In order to characterise our new biosensor module, we measured growth and fluorescence of *E. coli* harbouring pNarsenic in the presence of multiple naringenin concentrations and a wide gradient of NaAsO_2_. The dose-response curve measured in the absence of externally applied naringenin appeared entirely flat (Figure 1B). When 3-25 mg/L naringenin was added, the same biosensor produced a sigmoidal dose-response curve in response to arsenic. These results show that ArsR endogenously expressed by *E. coli* or originating from leaky expression from the P_*fdeAR*_ promoter was not present at a sufficient level to give rise to differential *mkate2* expression.

The response curves measured in the presence of 3-25 mg/L naringenin were fitted to a Hill function to extract quantitative parameters that enable comparison of biosensor performance (Table S2). Figure 1C shows that the basal output *b* decreased with increasing naringenin concentration. From equation (3), the dynamic range *µ* increased as a result (Figure 1D). Concurrently, the sensing threshold *θ* shifted to higher arsenic concentrations (Figure 1E). These trends can be explained by the mechanism of ArsR regulation. At higher naringenin concentrations, more ArsR dimers accumulate in the cell. This reduces *mkate2* expression in the absence of arsenic, thereby lowering the basal output. At the same time, higher arsenic concentrations are required to displace ArsR from the TFBS, as a result of which the threshold increases. These results highlight the tunability of the biosensor, making it adaptable for applications requiring different sensitivities.

The response curves measured in the presence of 6.5 and 10 mg/L naringenin overlapped to a large extent (Figure 1B). In addition, their values for *b, a* and *θ* did not differ significantly (Figure 1C & 1E and Table S2). These results suggest that the biosensor is not sensitive to this small increase in naringenin concentration.

The operational range of a biosensor, the span of effector concentrations that yield distinguishable output signals, is a key determinant of its practical applicability. At 3 mg/L naringenin, the operational range was 0.09 µM to 4.73 µM (Figure 1F), thus achieving sensitivity below the World Health Organisation limit for drinking water (0.133 µM) [16]. Upon increasing the naringenin concentration to 6.5 mg/L, the operational range shifted towards higher arsenic concentrations while maintaining approximately the same width on a logarithmic scale. At higher naringenin concentrations, however, the operational range constricted, with an approximately constant upper bound around 30 µM. This result suggests a practical limit for dose-response curve engineering by tuning transcription factor concentration alone. Future research should investigate whether such limits apply to other biosensors as well.

Finally, the noise level of a biosensor is an important factor to consider for monitoring applications [10]. pNarsenic exhibits relatively low noise levels between 15% and 32% (for 3 and 10 mg/L naringenin, respectively; Table S2), demonstrating its reliability.

### 3.3 Comparison with a phenomenological model for biosensor response

Because of the high quality of our data and fits, we were able to use them to extend the phenomenological model of Mannan *et al*. to the TF expression level, which is captured by the parameter *a*_1_ [8]. For low *a*_1_ values, corresponding to low TF expression levels, the model produced a high effector-independent response, in agreement with our results in the absence of naringenin (Figures 1B and 2). For increasing *a*_1_ values, the basal output decreased and the threshold and dynamic range increased, which is also supported by our data for pNarsenic. However, the maximum output *a* + *b* was independent of *a*_1_ (see Supplementary Information), which is in contrast with our data and data from a previous study [7], where the maximum output decreased markedly for higher-level TF expression. This observation probably results from incomplete derepression at high TF densities, a feature not accounted for by the model.

**Figure 2:**
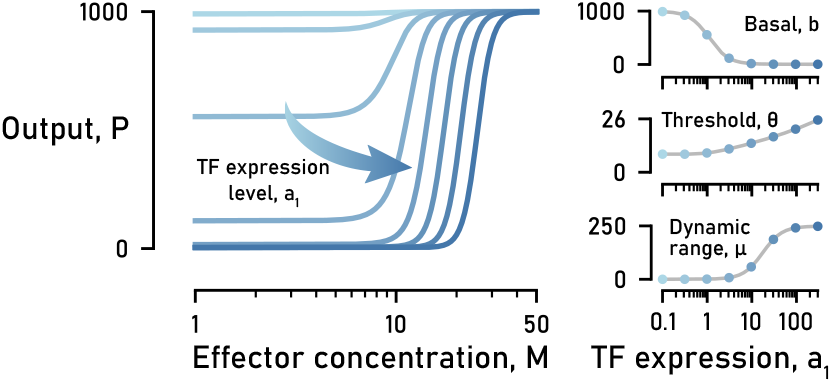
The phenomenological model of Mannan *et al*. [8] predicts that increasing the TF expression level (parameter *a*_1_) leads to decreasing basal output *b* and increasing threshold *θ* and dynamic range *µ*. Plots show simulations of the dose-response curve for eight values of *a*_1_ and the predicted behaviour of *b, θ* and *µ*. Simulations were made by setting *b*_1_ = 0.01, *b*_2_ = 4, *a*_2_ = 996, *K*_1_ = 0.1, *K*_2_ = 0.9, *n*_1_ = 6, *n*_2_ = 2 in equations (6) to (8) and (10) to (11) with *a*_1_ values spanning the range *a*_1_ = 0.1 to *a*_1_ = 300.

As shown in the Supplementary Information, the basal output and dynamic range are monotonic and converge to an asymptotic value for *a*_1_ → +∞, but this was not true for the threshold. Although we were not able to compute 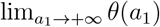 symbolically due to its complexity, simulations across a wide parameter scan suggest that the threshold went to infinity for *a*_1_ → +∞. This was expected as the model does not consider a limit for total intracellular protein. Besides, depending on the parameter values, *θ*(*a*_1_) could reach a minimum value (Figure S2), implying that further decreasing the transcription factor concentration increases the threshold. We were not able to attach a biological explanation to this phenomenon; it could be a mathematical artefact stemming from the nonlinear dependence of *f*_2_ on *f*_1_ in equation (6).

## Author information

### Author contribution

**Vincent Crabbe:** Conceptualization, Methodology, Software, Formal analysis, Investigation, Writing - Original Draft, Writing - Review & Editing, Visualization. **Ezgi Unal:** Investigation. **Stijn De Graeve:** Investigation. **Daniel G. Giraldez:** Funding acquisition. **Tom Peeters:** Funding acquisition. **Sophie de Buyl:** Methodology. **Eve-line Peeters:** Supervision, Funding acquisition. **Indra Bervoets:** Conceptualization, Methodology, Supervision.

### Note

No potential conflict of interest was reported by the authors.

## Acknowledgements

We thank Karl Jonckheere for technical assistance. The authors acknowledge the use of generative artificial intelligence for improving the clarity and style of this manuscript and for writing code. All conceptual, analytical, and interpretative aspects of the work were conducted by the authors.

## References

[1] Jan Roelof van der Meer and Shimshon Belkin. Where microbiology meets microengineering: Design and applications of reporter bacteria. Nature Reviews Microbiology, 8:511–522, 2010.

[2] Changjiang Liu, Huan Yu, Baocai Zhang, Shilin Liu, Chen-guang Liu, Feng Li, and Hao Song. Engineering whole-cell microbial biosensors: Design principles and applications in monitoring and treatment of heavy metals and organic pollutants. Biotechnology Advances, 60:108019, 2022.

[3] Joel Podgorski and Michael Berg. Global threat of arsenic in groundwater. Science, 368(6493):845–850, 2020.

[4] Raul Fernandez-López, Raul Ruiz, Fernando de la Cruz, and Gabriel Moncalián. Transcription factor-based biosensors enlightened by the analyte. Frontiers in Microbiology, 6:648, 2015.

[5] Jianhu Wu and Barry P. Rosen. Metalloregulated expression of the ars operon. Journal of Biological Chemistry, 268(1):52–58, 1993.

[6] Arthur Carlin, Weiping Shi, Saibal Dey, and Barry P. Rosen. The ars operon of escherichia coli confers arsenical and antimonial resistance. Journal of Bacteriology, 177(4):981–986, 1995.

[7] Baojun Wang, Mauricio Barahona, and Martin Buck. Amplification of small molecule-inducible gene expression via tuning of intracellular receptor densities. Nucleic Acids Research, 43(3):1955–1964, 2015.

[8] Ahmad A. Mannan, Di Liu, Fuzhong Zhang, and Diego A. Oyarzún. Fundamental design principles for transcription-factor-based metabolite biosensors. ACS Synthetic Biology, 6(10):1851–1859, 2017. doi: 10.1021/acssynbio.7b00172.

[9] Davide Merulla, Vassily Hatzimanikatis, and Jan Roelof van der Meer. Tunable reporter signal production in feedback-uncoupled arsenic bioreporters. Microbial Biotechnology, 6(5):503–514, 2013.

[10] Brecht De Paepe, Jo Maertens, Bartel Vanholme, and Marjan De Mey. Modularization and response curve engineering of a naringenin-responsive transcriptional biosensor. ACS Synthetic Biology, 7(5):1303–1314, 2018.

[11] Amber J. Bernauw, Vincent Crabbe, Fraukje Ryssegem, Ronnie Willaert, Indra Bervoets, and Eveline Peeters. Molecular mechanisms of regulation by a β-alanine-responsive lrp-type transcription factor from acidianus hospitalis. MicrobiologyOpen, 12(3):e1356, 2023.

[12] A. M. Marin, E. M. Souza, F. O. Pedrosa, L. M. Souza, G. L. Sassaki, V. A. Baura, M. G. Yates, R. Wassem, and R. A. Monteiro. Naringenin degradation by the endophytic diazotroph herbaspirillum seropedicae smr1. Microbiology, 159:167–175, 2013.

[13] Fernanda Miyuki Kashiwagi, Brenno Wendler Miranda, Emanuel Maltempi De Souza, and Marcelo Müller-Santos. The naringenin-dependent regulator fder can be applied as a nimply gate controlled by naringenin and arabinose. Synthetic Biology, 9(1), 2024.

[14] Dmitry Shcherbo, Christopher S. Murphy, Galina V. Ermakova, Elena A. Solovieva, Tatiana V. Chepurnykh, Aleksandr S. Shcheglov, Vladislav V. Verkhusha, Vladimir Z. Pletnev, Kristin L. Hazelwood, Patrick M. Roche, Sergey Lukyanov, Andrey G. Zaraisky, Michael W. Davidson, and Dmitriy M. Chudakov. Far-red fluorescent tags for protein imaging in living tissues. Biochemical Journal, 418(3):567–574, 2009.

[15] Sheng-Yan Chen, Yan Zhang, Renjie Li, Baojun Wang, and Bang-Ce Ye. De novo design of the arsr regulated pars promoter enables a highly sensitive whole-cell biosensor for arsenic contamination. Analytical Chemistry, 94(20):7210–7218, 2022. doi: 10.1021/acs.analchem.2c00055.

[16] Guidelines for drinking-water quality: fourth edition incorporating the first and second addenda. World Health Organization, Geneva, 2022.

