## Supplemental information for "Development and characterisation of pNarsenic: a naringenin-inducible biosensor for arsenic in *Escherichia coli*"

#### 1 Supplementary methods

##### 1.1 Strains and growth conditions

*Escherichia coli* MG1655 was used for all experiments. For plasmid construction, *E. coli* was grown in liquid lysogeny broth (LB) or solid LB (1.5% agar). For reporter gene assays, MOPS EZ Rich Defined Medium with 0.2% glucose was used as growth medium. Where appropriate, growth media were supplemented with 60 µg/mL kanamycin.

##### 1.2 Plasmid construction

Plasmids were constructed using NEBuilder® HiFi DNA Assembly (New England Biolabs). DNA fragments were purchased from Twist Bioscience, and DNA oligos were purchased from Sigma-Aldrich. KAPA HiFi DNA polymerase (Roche) was used for DNA amplification. The DNA sequences of the constructed plasmids were verified using whole-plasmid sequencing service (Eurofins Genomics). This revealed that the sequence of the plasmids were correct, but concatemers were present in the plasmid preparations.

#### 1.3 Reporter gene assays and data-analysis

The `model` function of the Python package LMFIT was used for fitting, weighting the residuals by the inverse of the mean fluorescence at the corresponding $NaAsO_2$ concentration. 95% confidence intervals of the regression parameters were determined using the `conf_interval` function.

A general formula for the operational range was derived by assuming that a biosensor cannot be used to translate a given fluorescent output to a specific effector concentration when the fluorescent output is too close to the basal output $b$ or the maximum output $a+b$. More formally, we defined the operational range as the range of arsenic concentrations where P([As]) satisfies:

$$b + f \cdot a \ \leq \ P([\mathrm{As}]) \ \leq \ b + (1-f) \cdot a$$

with $f$ the fraction of $a$ that we should not consider on both sides of the response curve. Substituting (2) into this equation and solving for [As], we obtain equation (4):

$$\mathrm{OR} = \left[ \sqrt[n]{\frac{f}{1-f}} \cdot \theta, \ \sqrt[n]{\frac{1-f}{f}} \cdot \theta \right]$$

#### 1.4 Analysis of the phenomenological model of Mannan *et al.* (2017)

Table S1 shows the definition of the model parameters of the phenomenological model of Mannan *et al.* (2017).

Table S1: Definition of the model parameters of the phenomenological model of Mannan *et al.* (2017). Abbreviations: TF, transcription factor; TFBS, transcription factor binding site.

| Parameter | Definition |
|---|---|
| $b_1$ | Basal level of TF activity |
| $b_2$ | Basal level of promoter expression |
| $a_1$ | Maximum increase in TF activity |
| $a_2$ | Maximum increase in promoter expression |
| $K_1$ | Effector-TF affinity |
| $K_2$ | TF-TFBS affinity |
| $n_1$ | Sensitivity of effector-TF binding |
| $n_2$ | Sensitivity of TF-TFBS binding |

The equations used to calculate the basal output, maximum increase in output, threshold, and dynamic range were taken from Mannan *et al.* (2017):

$$b = b_2 + \frac{a_2}{1 + (K_2(b_1 + a_1))^{n_2}} \tag{8}$$

$$a = a_2 \cdot \frac{\left((b_1 + a_1)^{n_2} - b_1^{n_2}\right) K_2^{n_2}}{\left(1 + (K_2\,(b_1 + a_1))^{n_2}\right)\left(1 + (b_1 K_2)^{n_2}\right)} \tag{9}$$

$$\theta = \frac{1}{K_1} \sqrt[n_1]{\frac{a_1 K_2}{\sqrt[n_2]{A - 1} - b_1 K_2} - 1} \tag{10}$$

$$\mu = \mu_2 \cdot \frac{((b_1 + a_1)^{n_2} - b_1^{n_2}) K_2^{n_2}}{(1 + (K_2(b_1 + a_1))^{n_2} + \mu_2) \cdot (1 + (b_1 K_2)^{n_2})} \tag{11}$$

with

$$A = 2 \cdot \frac{(1 + (K_2(b_1 + a_1))^{n_2}) \cdot (1 + (b_1 K_2)^{n_2})}{2 + ((b_1 + a_1)^{n_2} + b_1^{n_2}) \cdot K_2^{n_2}}$$

$$\mu_2 = \frac{a_2}{b_2}$$

We treat all parameters as constants except for $a_1$. As such, we consider $b(a_1)$, $a(a_1)$, $\theta(a_1)$ and $\mu(a_1)$ as functions of $a_1$.

### 2 Supplementary results

#### 2.1 *In vivo* characterisation of pNarsenic

This section lists the dose-response parameters obtained for pNarsenic (Table S2) and shows the dose-response curves from Figure 1B in more detail (Figure S1).

#### 2.2 Comparison with a phenomenological model for biosensor response

This section covers the results regarding the phenomenological model from Mannan *et al.* (2017) that support our conclusions in the main text. From equations (8) and (9), we find that:

$$a + b = b_2 + \frac{a_2}{1 + (b_1 K_2)^{n_2}} = \lim_{M \to +\infty} P(M)$$

from which we conclude that the maximum output $a + b$ is not a function of $a_1$.

To derive the global behaviour of the the basal output and the dynamic range with respect to $a_1$, we computed the derivatives of equations (8) and (11) with the help of SymPy:

$$\frac{\mathrm{d}b}{\mathrm{d}a_1} = -\frac{a_2 n_2 K_2^{n_2} (b_1 + a_1)^{n_2 - 1}}{\left[1 + (K_2\,(b_1 + a_1))^{n_2}\right]^2} \tag{12}$$

$$\frac{\mathrm{d}\mu}{\mathrm{d}a_1} = \left(1 + \frac{\mu_2}{1 + (b_1 K_2)^{n_2}}\right) \cdot \frac{\mu_2\, n_2\, K_2^{n_2}\, (b_1 + a_1)^{n_2 - 1}}{\left[1 + \left(K_2\,(b_1 + a_1)\right)^{n_2} + \mu_2\right]^2} \tag{13}$$

Table S2: Dose-response parameters for the pNarsenic biosensor in the presence of the indicated naringenin concentration. Reported values for the basal output ($b$), maximum increase in output ($a$), threshold ($\theta$), and Hill coefficient ($n$) are the best-fit values along with their 95% confidence interval between parentheses. The dynamic range ($\mu$), operational range, and Noise parameter were computed from equations (3) to (5).

| Naringenin (mg/L) | $b$ (a.u.) | $a$ (a.u.) | $\theta$ (µM) | $n$ | $a+b$ | $\mu$ | Operational range (µM) | Noise (%) |
|---|---|---|---|---|---|---|---|---|
| 3 | 785 (660 - 901) | 5353 (4869 - 5862) | 0.64 (0.47 - 0.91) | 1.1 (0.88 - 1.38) | 6138 | 6.8 | [0.09, 4.73] | 15 |
| 6.5 | 139 (103 - 173) | 7371 (6296 - 8582) | 3.29 (2.04 - 5.74) | 0.96 (0.82 - 1.15) | 7509 | 53 | [0.33, 32.29] | 25 |
| 10 | 147 (105 - 187) | 6625 (5346 - 8069) | 3.27 (1.98 - 6.07) | 1.3 (1.03 - 1.68) | 6772 | 45 | [0.60, 17.85] | 32 |
| 25 | 52 (44 - 60) | 4550 (3898 - 5263) | 6.68 (4.92 - 9.55) | 1.53 (1.32 - 1.8) | 4602 | 87 | [1.59, 28.06] | 18 |

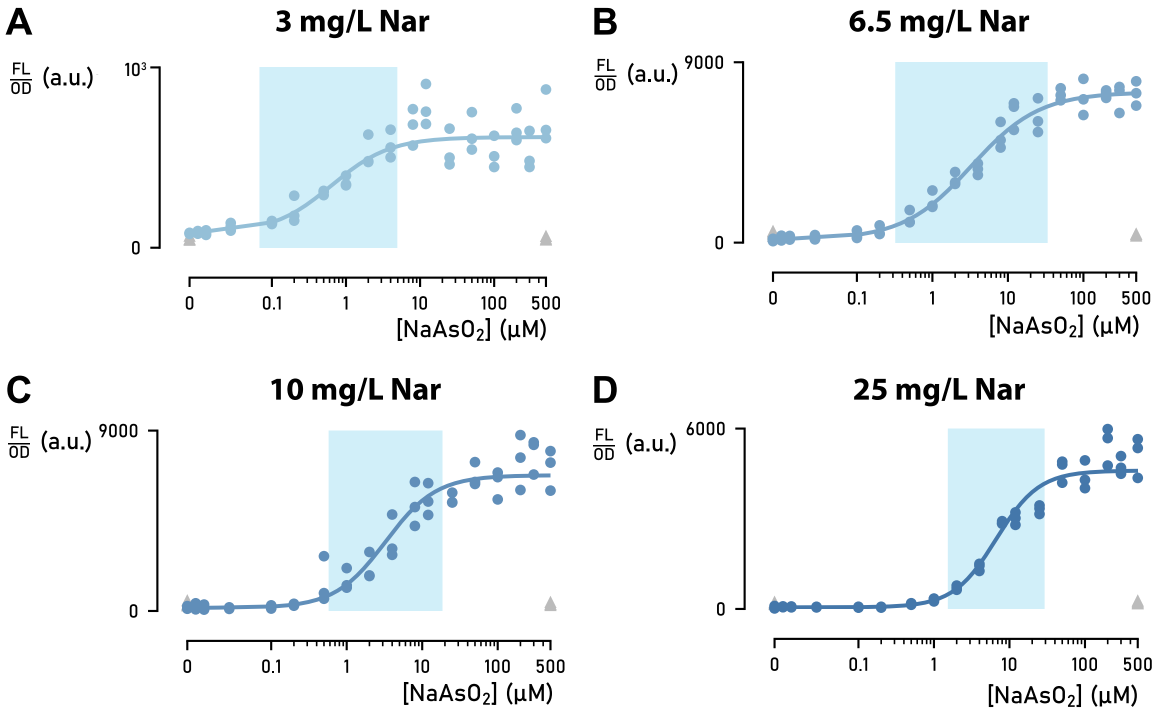


Figure S1: Dose-response curves of the pNarsenic biosensor and their best-fit Hill functions. Subplots show the relative fluoresence output $\frac{\mathrm{FL}}{\mathrm{OD}}$ in the presence of 3 mg/L (A), 6.5 mg/L (B), 10 mg/L (C), and 25 mg/L (D) naringenin for the pNarsenic biosensor as blue dots. Grey triangles show the $\frac{\mathrm{FL}}{\mathrm{OD}}$ for a control plasmid lacking the $\mathrm{P}_{arsOC2}$ promoter upstream of *mkate2*. The light-blue shaded box indicates the operational range as computed by (4). The x-axis scale is linear between 0 µM and 0.1 µM and logarithmic between 0.1 µM and 500 µM.

As all parameters are strictly positive, it follows from equations (12) and (13) that $b$ is monotonically decreasing and $\mu$ is monotonically increasing. In addition, $b$ and $\mu$ evolve to an asymptotic value for $a_1 \to \infty$:

$$\lim_{a_1 \to +\infty} b = b_2 \tag{14}$$

$$\lim_{a_1 \to +\infty} \mu = \frac{\mu_2}{1 + (b_1 K_2)^{n_2}} \tag{15}$$

For $\theta$, we obtained the following derivative:

$$\frac{\mathrm{d}\theta}{\mathrm{d}a_1} = \frac{1}{K_1 n_1} u^{\frac{1}{n_1}-1} \cdot \left[ \frac{K_2}{\sqrt[n_2]{A-1} - b_1 K_2} - \frac{a_1 K_2}{\left(\sqrt[n_2]{A-1} - b_1 K_2\right)^2} \cdot \frac{1}{n_2} (A-1)^{\frac{1}{n_2}-1} \frac{\mathrm{d}A}{\mathrm{d}a_1} \right] \tag{16}$$

with

$$u = \frac{a_1 K_2}{\sqrt[n_2]{A-1} - b_1 K_2} - 1$$

$$\frac{\mathrm{d}A}{\mathrm{d}a_1} = 2 \cdot \frac{n_2 K_2^{n_2} \left(b_1 + a_1\right)^{n_2 - 1} \cdot \left(1 + \left(b_1 K_2\right)^{n_2}\right)}{2 + \left(\left(b_1 + a_1\right)^{n_2} + b_1^{n_2}\right) \cdot K_2^{n_2}}$$
$$- 2 \cdot \frac{n_2 K_2^{n_2} \left(b_1 + a_1\right)^{n_2 - 1} \left(1 + \left(K_2 \left(b_1 + a_1\right)\right)^{n_2}\right) \left(1 + \left(b_1 K_2\right)^{n_2}\right)}{\left(2 + \left(\left(b_1 + a_1\right)^{n_2} + b_1^{n_2}\right) \cdot K_2^{n_2}\right)^2}$$

We could not solve equation (16) to an explicit expression for $a_1$, but one finds after a little algebra that setting (16) to zero is equivalent to

$$\sqrt[n_2]{A-1} - b_1 K_2 - \frac{a_1}{n_2} \cdot (A-1)^{\frac{1}{n_2} - 1} \frac{\mathrm{d}A}{\mathrm{d}a_1} = 0 \qquad (17)$$

Next, we sampled the parameter space using latin hypercube sampling. For each random parameter set, we substituted the values of $b_1$, $b_2$, $a_2$, $K_1$, $K_2$, $n_1$, and $n_2$ into equation (17) and solved numerically to $a_1$ using the `nsolve` function of SymPy with the sampled value of $a_1$ as initial guess. For 164 out of 500 parameter sets, we found a numerical solution, in each case indicating a minimum value for $\theta(a_1)$. Figure S2 shows one example of such a minimum; similar simulations for the 163 other parameter sets can be consulted in the Supplementary Data. Numeric analysis showed that $\theta$ goes to infinity for $a_1 \to +\infty$. We conclude that a minimum to $\theta(a_1)$ can exist depending on the parameter values.

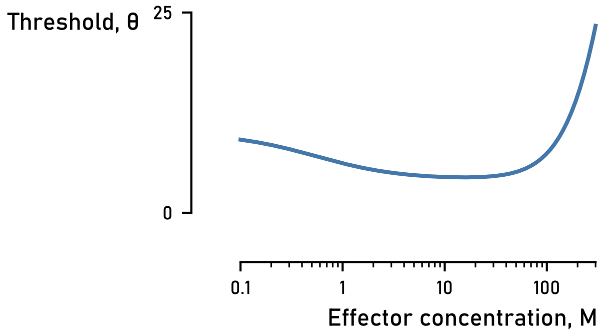


Figure S2: Simulation showing a minimum value for the threshold. The simulation was made by setting $b_1 = 0.5$, $b_2 = 10$, $a_2 = 9000$, $K_1 = 0.1$, $K_2 = 0.01$, $n_1 = 1$, $n_2 = 2$ in equation (10) with $a_1$ values spanning the range $a_1 = 0.1$ to $a_1 = 300$.
